# The effect of glucose restriction on cancer cell contractility: A threshold response in U-87 glioma

**DOI:** 10.1101/2024.09.05.611526

**Authors:** Albert Kong, Alessia Pallaoro, Donald Yapp, Gwynn J. Elfring, Mattia Bacca

## Abstract

Cells rely on contractility to proliferate, and cancerous ones exhibit an increased glucose dependence. It is therefore hypothesized that glucose restriction can mitigate cancer cell proliferation by ’stunting’ their contractility. However, glucose-restriction studies have mostly been based on experiments that have yielded conflicting results; some cells become less contractile under glucose-restriction, intuitively, while, others become surprisingly more contractile. Active mechanistic modeling may prove fruitful in resolving these conflicts. In this study, we develop a model for glucose-mediated cell contractility to capture the mechanical implications of glucose restriction. The model is calibrated on cell contraction data taken from 2D-cultured glioma cells, laying on a collagen substrate. The model predicts the existence of a critical level of glucose restriction that must be exceeded for contractility to be affected, and this is validated by our experiments. Our model provides an initial step toward a fundamental understanding of the metabolic implications of cell contractility, particularly in the context of glucose restriction: an essential step in cancer studies.

**significance:** This study advances our understanding of how glucose restriction affects cancer cell contractility, an essential factor in cell proliferation. Our findings reveal that cells require severe glucose deprivation before exhibiting reduced contractility, highlighting a threshold response. This indicates that the cytoskeleton, a key structural component, remains active until a significant reduction in energy supply forces the cell into a lower energy state. These insights provide critical knowledge about the metabolic hierarchy within cells, contributing to the broader study of cancer metabolism and potential therapeutic strategies aimed at disrupting cellular energy pathways.

---

Cellular energy is stockpiled as adenosine triphosphate (ATP) and hydrolyzed to perform useful functions such as signaling, active transport, chemical synthesis, motility, and contractility (1). Cells can produce ATP in various ways, often via breaking down glucose. When oxygen is abundant this is done through aerobic glycolysis, but in the absence of oxygen (hypoxia), the less efficient (fewer ATP molecules per glucose) anaerobic glycolysis pathway is used (2). In a phenomenon known as the Warburg effect, cancer cells produce ATP preferentially through anaerobic glycolysis, irrespective of their oxygen content (3, 4). Consequently, cancer cells require substantially more glucose to remain functional, making them more sensitive to glucose availability.

In an attempt to exploit the Warburg effect, studies have employed glucose restriction to investigate potential nutritional or dietary approaches for cancer therapy (5, 6). These studies have predominantly been experimental, and primarily focused on answering questions about physiological, biochemical, and or rate-of-mortality-related changes in cancer cells (7–12). Furthermore, these studies have yielded conflicting results (13). Intuitively, some cells become less active and proliferative under glucose restriction, due to a higher requirement for energy efficiency, while others become surprisingly more active (13). This unexplained phenomenon could be due to an innate attempt of cells to spread toward regions of nutrient abundance. Cancer cells rely on contractility to proliferate (14, 15), thus, a mechanistic model that relates contractility to glucose availability may help to shed light on the conflicting experimental results observed in the literature. In this paper, we develop a simple mechanistic model for cell contraction as a function of glucose restriction. Our model results predict a critical threshold in glucose restriction that must be trespassed to reduce cell contractility, in agreement with our experiments. Thus, mild levels of glucose restrictions leave cell contractility unaffected or even enhanced. Our observation might resolve the current contradictions in the literature, highlighting the [1] importance of quantitative assessments in glucose restriction studies.

## Cell Contraction Model

Our model idealizes cells as an active contractile mechanism that responds to external stimuli (Figure.1). In its rest state, the cell spreads over and anchors on to a substrate via clusters of adhesive or receptor-ligand binding molecules called focal adhesions (FAs). The cell’s primary contractile structures are stress fibres (SF); bundles of actin filaments and myosin motors. The cross-bridge cycling of myosin motors in SFs generates tension, which is resisted by FAs that terminate at either end of SFs. Cells contract by remodelling both FAs and SFs: FAs slide via protein treadmilling while SFs shorten via protein dissociation. Both must occur simultaneously for contraction to occur. In this section, we detail how the remodelling of FAs and SFs is mediated by the active mechanotransduction of biochemical energy (ATP hydrolysis).

### Focal adhesion remodeling

If the tension in the stress fibers *T*_*SF*_ is sufficiently large, the focal adhesion sites will slide and the stress fibers will shorten, but how FAs transition between their static and sliding states is still poorly understood (16). FA sliding occurs via traction-mediated protein tread-milling, with FA subunits dissociating from the peripheral edge while associating at the central edge (Figure 1a). Interestingly, during sliding, FAs have been observed to maintain a constant area 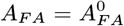 (when viewed orthogonal to the substrate plane) (16–18). Further, the rate at which proteins turnover in FA complexes - and consequently the FA sliding velocity - was found to be positively affected by the shear stress sustained by the FA complex *σ*_*FA*_ (19). Note that *σ*_*FA*_ must balance the tension in SFs

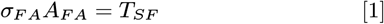

**Fig. 1.**
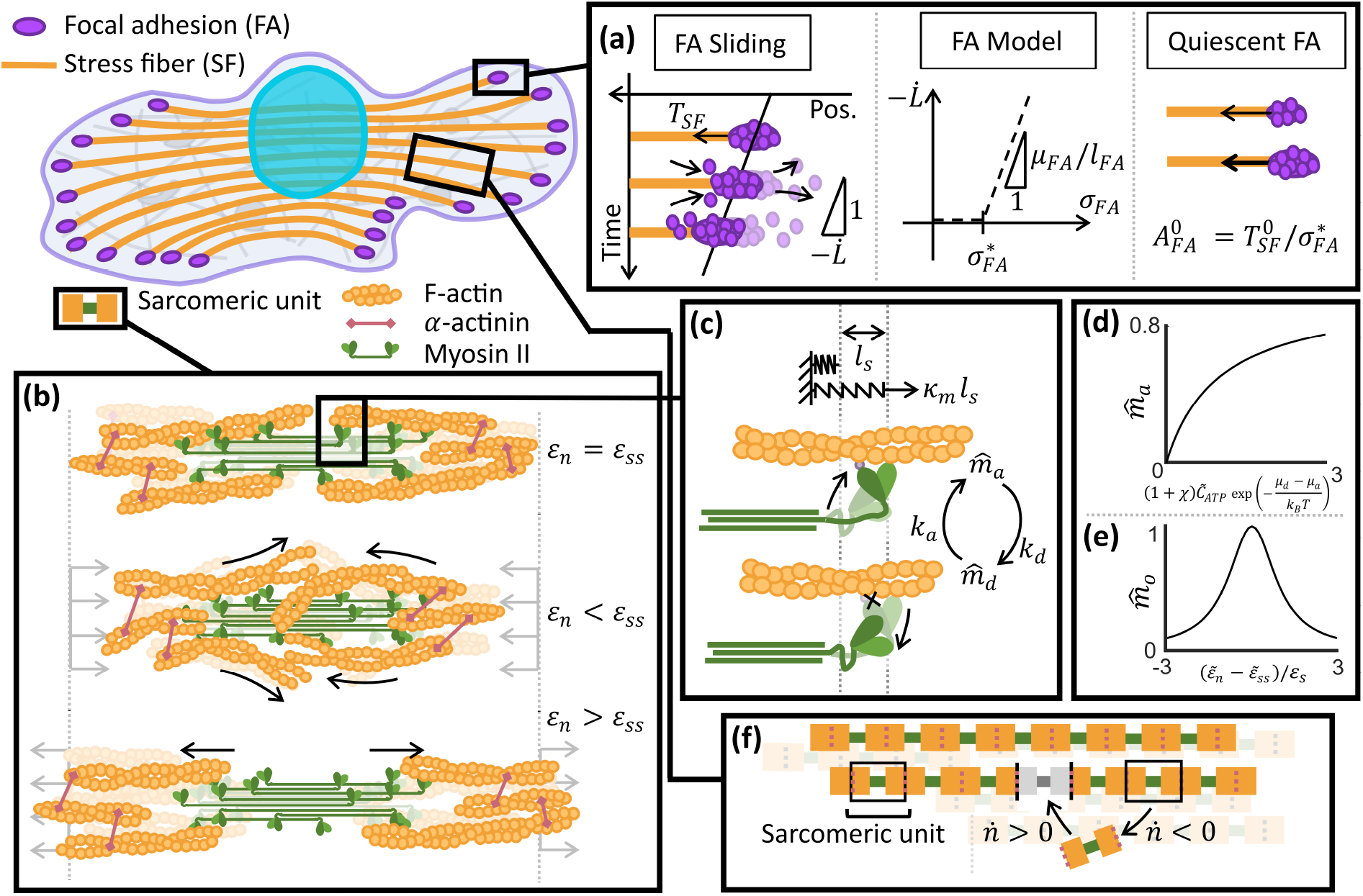
Overview of the cell contraction model. Cell contraction is driven by a network of contractile stress fibers (SFs): filamentous cytoskeletal structures that are assembled from serially repeating sarcomeric units. We assume that SFs are predominantly aligned along the cell length (*L*), with homogeneously evolving properties. SFs are anchored at their ends to focal adhesion sites (FAs). (a) FAs are layered, trans-membrane protein complexes that anchor SFs to the substrate at their ends. When cells contract, FA proteins dissociate from their peripheral edge and associate at the central edge. The net effect is an apparent sliding whereby FA complexes maintain their size and shape. We model the bi-stability of FAs using a linear visco-plastic model. Below a critical shear stress 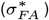 FAs remain stationary, while above it, it slides at a rate proportional to the excess shear stress 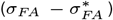. In quiescent cells, the size/area of FAs during sliding 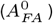 scale with the sustained SF tension initially 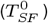. (b) Illustration of the structure of a sarcomeric unit, with emphasis on the organization of its components: F-actin, *α*-actinin, and Myosin II bipolar filaments, and how it changes with strain (*ε*_*n*_). When strained away from the steady-state value (*ε*_*ss*_), the overlap between actin and myosin diminishes, which in turn hampers the ability for the SF to generate tension. (c) Tension is generated by cross-bridge cycling myosin heads. We model cross-bridge cycling with a simple two-state model where the myosin heads are either attached or detached. The fraction of heads in either state is given by 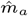 and 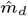 respectively. In the attached state, the elastic necks of the myosin are stretched by *l*_*s*_, which generates a force of *κ*_*m*_*l*_*s*_ in the SFs, with *κ*_*m*_ the stiffness of the necks. In the detached state, the heads are separated from actin and do not contribute to tension generation. We model the transition of the myosin head fractions via simple reaction kinetics, with attachment and detachment rates given by *k*_*a*_ and *k*_*d*_ respectively. (d) We model the effects of strain (*ε*_*n*_ − *ε*_*ss*_) on the fraction of actively cross-bridge cycling myosin heads 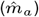 using a Cauchy distribution with parameter *ε*_*s*_. (e) The fraction of attached myosin heads (*m*_*a*_) as a function of the ATP concentration 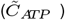, contractile signal (*χ*), and difference between detached (*µ*_*d*_) and attached (*µ*_*a*_) head enthalpies. (f) SFs can lengthen or shorten by increasing 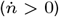 or decreasing g 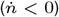 their number of sarcomeric units (*n*), respectively, but we only consider the latter in our contractile model.

We elect to minimally model the traction-dependent bistability of FAs using a yield stress fluid model. We assume that below a critical FA stress 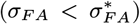, FAs remain stationary, but above it, protein turnover is enhanced, and the FAs slide following the relation

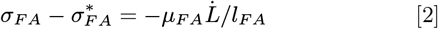

where *µ*_*FA*_ is a characteristic viscosity, *L* is the SF length (equal to the distance between two opposite FAs), and *l*_*FA*_ is a characteristic length scale for the viscous dissipation (see SI). Here, we make explicit that the FA sliding velocity is equal to the SF shortening velocity 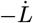. Both *µ*_*F A*_ and 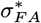 depend on the composition and concentration of FA constituents in the cytosol, which we assume to be constant for a given cell phenotype.

In quiescent cells, *A*_*FA*_ scales linearly with the tension exerted by the attached SFs (20–24). By assuming that prior to signaling, quiescent cells are at the cusp of contracting 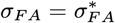 we find that 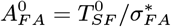 where 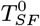 is the SF tension in a quiescent cell.

### Stress fiber tension generation

Cytoskeletal tension is generated via a complex molecular infrastructure called stress fiber, which base unit is the sarcomere. Figure 1b illustrates the structure of the sarcomeric units in detail. Here, Myosin II bipolar filaments are located centrally in sarcomeric units, surrounded at either end by antiparallel networks of *α*-actinin crosslinked actin filaments.

We adopt a two-state model for the cross-bridge cycle, where the myosin II heads are either attached or detached (Figure 1c), with fractions in either state given by 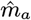 and 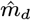 respectively. Attached myosin heads hydrolyze ATP to perform a power stroke, which stretches their elastic necks and generates tension in the SF. Detached heads, on the other hand, are separated from actin and do not contribute to tension generation (25).

The characteristic time for cross-bridge cycling (≈1 *ms*) is substantially shorter than cell contraction events (≈1000 *s*) (26–28). Consequently, the fraction of myosin heads in either state evolves effectively instantaneously (quasi-equilibrium), and independently from the contractile response. We assume that in the attached state, each myosin head contributes *κ*_*m*_*l*_*s*_ to the total SF tension, with *κ*_*m*_ the stiffness of the myosin neck and *l*_*s*_ their characteristic stretch (Hooke’s law). The total tension in a SF is then due to the total number of stretched myosin necks (in any unit)

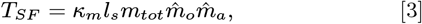

where *m*_*tot*_ the total number of myosin heads in a sarcomeric unit, which must be multiplied by the fraction of heads that are cross-bridge cycling 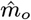 and the fraction of those that are attached 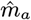.

For 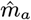 we model the myosin head transitions as a first-order reaction which, with the quasi-equilibrium assumption means that

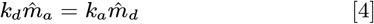

where *k*_*a*_ and *k*_*d*_ are the attachment and detachment rate coefficients respectively, modelled here with an Arrhenius-like law (29),

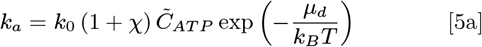

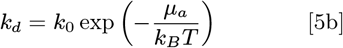

with *k*_0_ a nominal reaction rate constant; *χ* a dimensionless coefficient that provides the magnitude of the contractile signal; 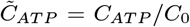 the dimensionless concentration of ATP (*C*_*ATP*_ is ATP concentration, and *C*_0_ a nominal concentration); *µ*_*a*_ and *µ*_*d*_ are the enthalpies of the attached and detached myosin heads, respectively; *k*_*B*_ is the Boltzmann constant; *T* is the system’s absolute temperature in Kelvin. By substituting Eq.5 into Eq.4, and the conservation of the total myosin head fractions 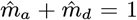, we arrive at the following equation, which gives the instantaneous fraction of attached myosin heads

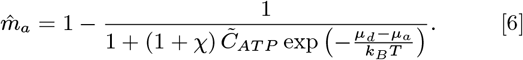

The dependence of the attached head fraction 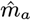 on the functional argument in Eq.6 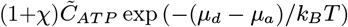 is illustrated in Figure 1d, where we see that heightened contractile signaling and ATP concentration lead to greater fractions of attached myosin heads, and subsequently, greater SF tension.

For 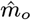 the relative motion between the myosin II bipolar filament and actin alters the strain, and in turn, the overlap between actin and myosin in the SF (Figure 1b). Stretching SFs from their steady-state configuration (with strain *ε*_*n*_ *> ε*_*ss*_) voids the SF center of actin, while compression (*ε*_*n*_ *< ε*_*ss*_) causes centrally located actin filaments to interfere (30, 31); both effects decrease the number of myosin heads available for cross-bridge cycling. Following (29) we approximate this effect by a Cauchy type distribution

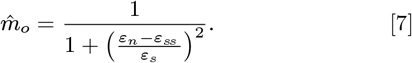

where the parameter *ε*_*s*_ controls the width of the bell-shaped distribution provided by Eq.7, and plotted in Figure 1e. *ε*_*s*_ depends on the structure of the functional units in the SF.

### Stress fiber remodeling

A stress fiber of length *L* consists of *n* sarcomeric units of length *l* serially connected such that *L* = *nl* as shown in Figure 1f. Generally, these variables vary with both location and orientation within the cell. However, SFs in contractile cells, particularly those cultured on stiff substrates, are often strongly aligned (32–34). We therefore simplify our analysis by assuming a perfectly aligned SF network. We further simplify by considering representative spatial averages of these variables, ultimately rendering our model as uniaxial and 0-dimensional.

Defining reference values *L*_*r*_ = *n*_*r*_*l*_*r*_ we may write in terms of strain relative to the reference

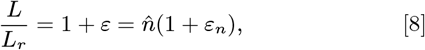

where *l/l*_*r*_ = 1 + *ε*_*n*_ and 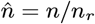. Now, a change in the SF length may be due to changes in the number of sarcomeric units or their length 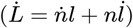, so we may write the rate of change of stress fiber length in terms of rates of strain as

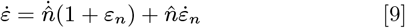

In our analysis, we assume that the cross-sectional area of the SFs bundle is constant. This is because the process of addition or subtraction of functional units along one fiber is much faster than the complete formation or dissociation of a fiber, required to remodel the cross-sectional area of the SF. To determine the strain *ε*_*n*_ in each sarcomeric unit, we follow a kinetic model shown in (35) to capture the stress fiber remodelling dynamics under strain. The complete SF kinetic model accounts for both lengthening, by the introduction of new sarcomeric units 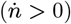, and shortening, by their removal 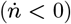. However, as we are concerned with contractile events alone, we consider only the latter in our analysis. When a sarcomeric unit is removed, the two bonds joining it to adjacent units must be broken, therefore the bond breaking rate is 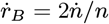. Removing sarcomeric units into the bulk results in the Newtonian dissipation law 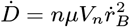, giving

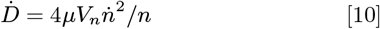

where *µ* is an effective viscosity and *V*_*n*_ is an effective volume of the dissipative process. The energy required to break these bonds originates from the reduction of SF configurational energy Ψ, such that

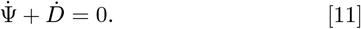

where Ψ = *nψ*, with *ψ* is the configurational internal energy of the unit sarcomere. The energy *ψ* (a type of aggregate bond potential of the sarcomeric unit) has minimimum *ψ*_0_ at *l*_*r*_, and we model the increase of this internal energy with strain with the power law

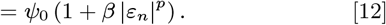

where *β* and *p* are model parameters. The rate of change of total configurational energy is then 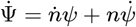, giving

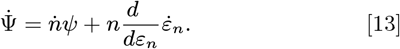

Upon substitution of Eq. (9) we obtain

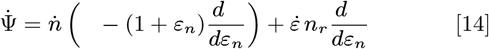

Because SF remodeling occurs at a faster rate than FA remodeling (due to the larger extension of FA complexes compared to the size of a single unit inside a SF), we can assume that 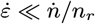, and thus neglect the second term on the right-hand side of Eq. (14). Equating now the dissipation in Eq. (11) and the free energy rate in Eq. (14) gives a differential equation for the number of sarcomeric units, or *ε*_*n*_ by the relationship in Eq. (8). Finally, by setting 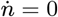, at a given macroscopic strain *ε*, we obtain an implicit equation for the steady-state strain *ε*_*ss*_

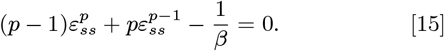

## Model Validation

Our model for ATP-mediated cell contraction is compared against contraction experiments. We subjected 2D cultured U-87 glioma cells to varying levels of glucose restriction and quantified changes in their contractility (Materials and Methods). We used UV and blue light stimulation (See SI) to trigger cell contraction during imaging and used image analysis to extract geometric contraction data against time.

### Experimental Results

In Figure 2a, we provide cell images obtained at the start (*t*_0_) and end (*t*_*f*_) of our imaging window (duration: 45 minutes). Qualitatively, the morphologies of the control (red), glucose-restricted (blue), and intermediate (green) cells were indistinguishable from one another at both timepoints.

**Fig. 2.**
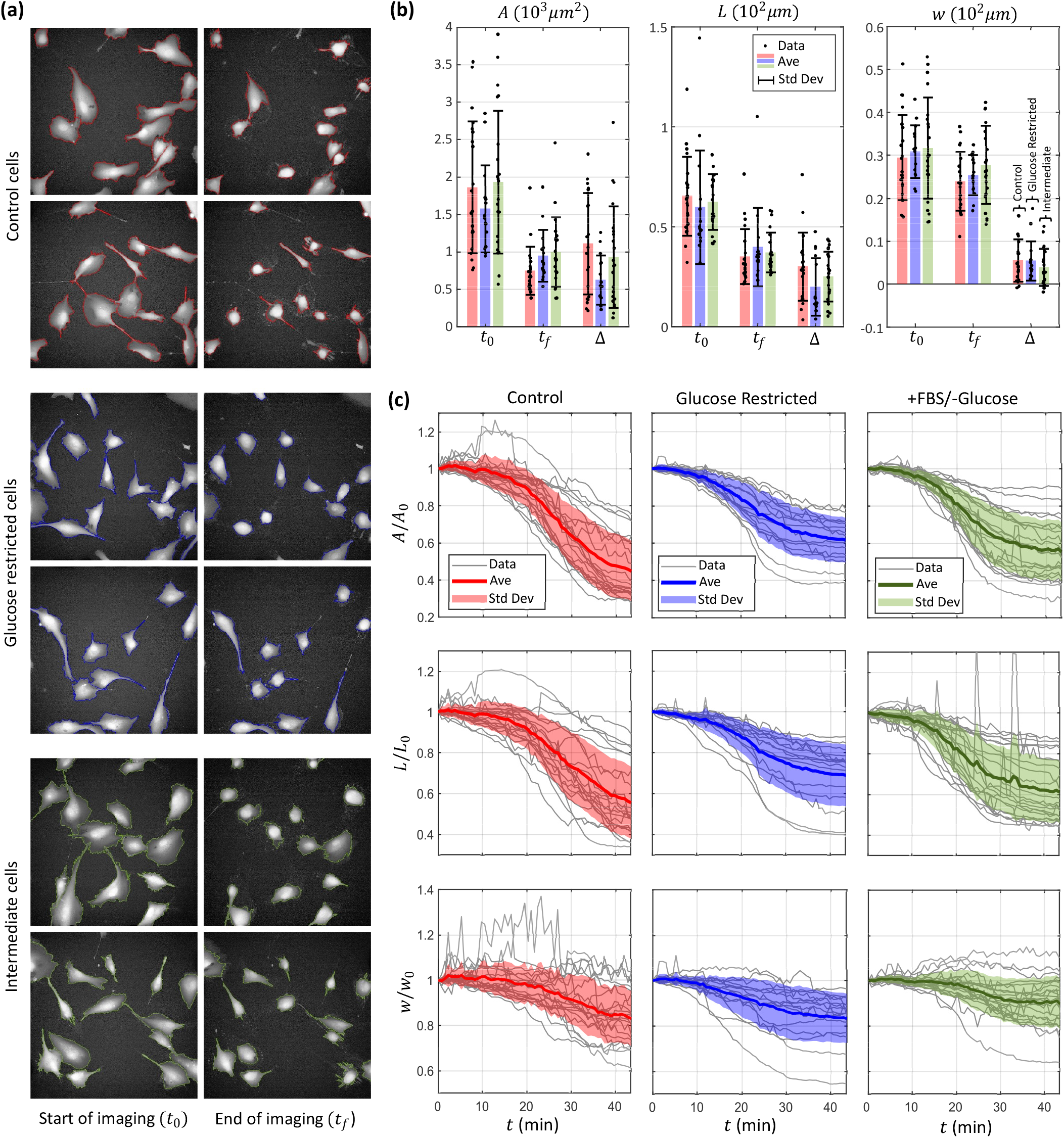
Summary of experimental results, data for control, glucose-restricted, and intermediate cells are color-coded red, blue, and green respectively. 21 control, 15 glucose-restricted, and 21 intermediate cells were studied. (a) Cell images at the start (*t*_0_) and end of imaging (*t*_*f*_), each square image is 333^2^ *µm*^2^ in area. (b) Bar graphs of the cell area (*A*), length (*L*), and width (*w*) at the start and end of imaging, as well as their difference (Δ). (c) Time series plots of the normalized cell area (*A/A*_0_), length (*L/L*_0_), and width (*w/w*_0_) with respect to their initial values (*A*_0_, *L*_0_, *w*_0_).

To facilitate quantitative analysis, we characterized the cells’ geometry by computing their area (*A*), length (*L*), and width (*w*) - with the cell length specified by the longer cell dimension initially (Materials and Methods). Figure 2b plots these quantities at the start and end of imaging, as well as their difference (Δ*A*, Δ*L*, and Δ*w*). Here, we confirm that there is no substantial difference in the start and end *A, L*, and *w* of our cells. We then normalized the cell areas, lengths, and widths by their starting values (*A*_0_, *L*_0_, *w*_0_ respectively), and plot their evolution over time in Figure 2c. By visualizing the contraction signatures, we find that they are sigmoidal in shape, and that, except for a few spreading control cells initially, all cells were contractile throughout. Furthermore, we find that by the end of the imaging window, glucose-restricted and intermediate cells have ceased to contract while several control cells remain contractile (seen in the non-zero contraction curve slopes at the end).

From Figure 2b, we find that the cells contracted minimally along their width (Δ*W* ≈ 45*µm*) while they contracted substantially more in their length (Δ*L* ≈ 250*µm*). Figure 2c demonstrates that this is also true for the normalized contractions. The cells contracted in their width (Δ*w/w*_0_) only by around 15% while they contracted in their length (Δ*L/L*_0_) by around 38% (more than a factor of 2 greater). This is to be expected because the cells were cultured on a stiff plastic substrate (Materials and Methods), and would have subsequently developed a strongly aligned stress fiber network along their length (their longer dimension) (32–34). This observation supports our model hypothesis of unidirectional contraction in cells, and we will only analyze contraction along the length of the cells when comparing our model with experiments.

### Model Validation

We first calibrate our model against control and glucose-restricted contraction data, as explained in the next sections, and then validate our predictions against experiments for intermediate glucose restriction (Figure 2c). Eq.14,2, and 3 constitute the main equation system of our model, respectively governing the evolution of SF kinetics (*n, ε*_*n*_) and tension generation (*T*_*SF*_). These are supplemented by Eq.12, 15, 7, 6, and 19. Missing from the model are definitions for the dimensionless ATP concentration 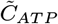 and contractile signaling *χ*, which we use for our calibration process.

### ATP Concentrations

Studies have estimated that as little as 20% of the total ATP production in cells is consumed by the actin cytoskeleton (36), and only a fraction of which is represented by SFs. Consequently, the functional form of 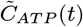 should have little direct relation to the observed contractile response. For simplicity we assume ATP consumption evolves linearly with time during the observed phenomenon, from which

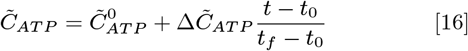

with 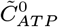 the normalized ATP concentration at the start and 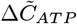 the change in normalized ATP concentration over the imaging window, going from *t*_0_ and *t*_*f*_.

By preparing parallel plates of control and glucose-restricted cells, and subjecting them to the same experimental procedures as the imaged cells in Figure 2a, we were able to measure cellular ATP content before and after imaging (see Materials and Methods). Figure 3a reports the ATP data, normalized against the ATP concentration in the control cells before imaging.

**Fig. 3.**
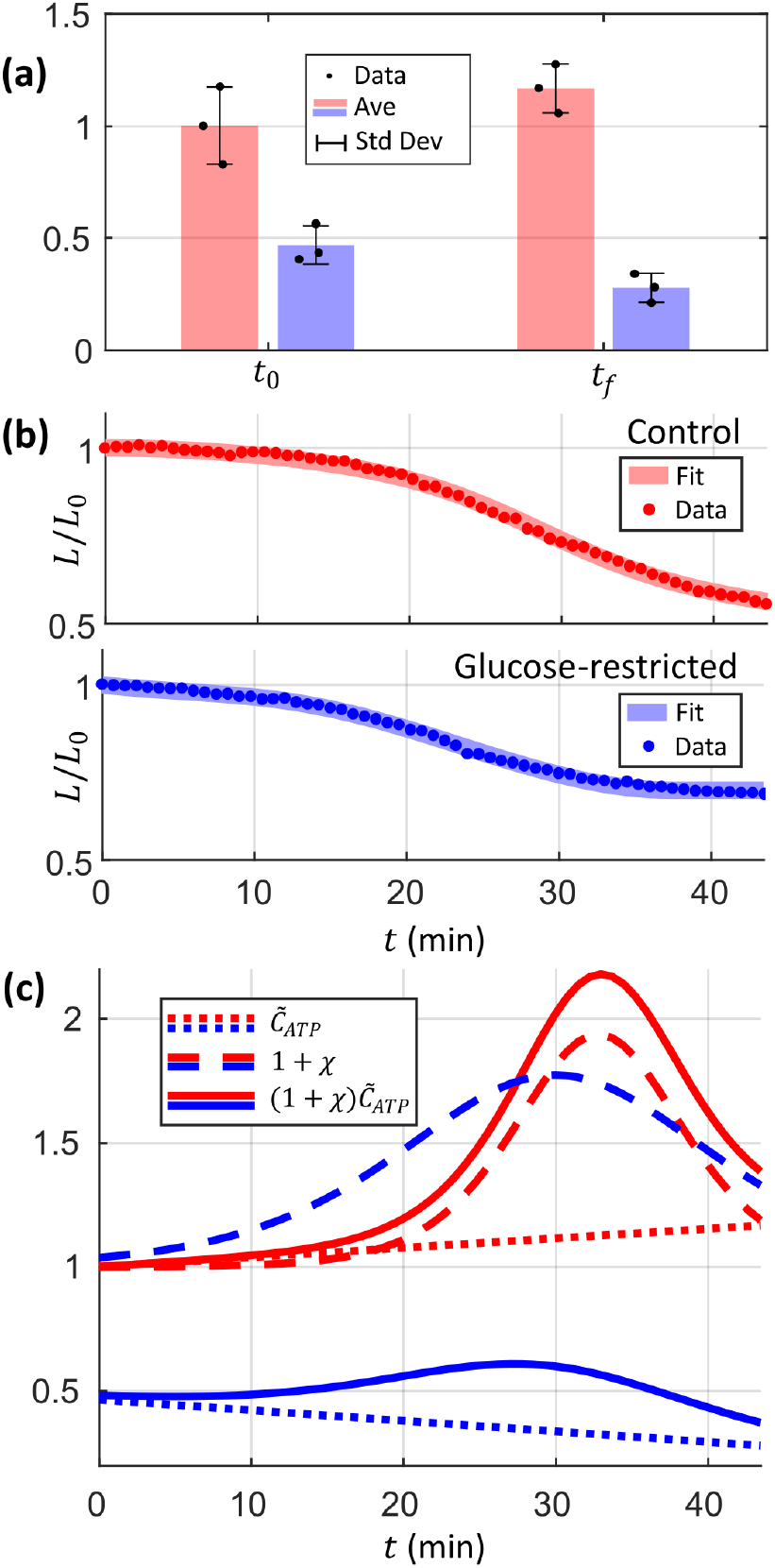
Results of fitting. (a) Cellular ATP concentration at the start and end for the control and glucose restricted cells, normalized against the average control value at the start 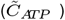. (b) The The control (*R*^2^ = 0.998) and glucose-restricted (*R*^2^ = 0.997) contraction signatures were captured well with our model. (c) The normalized ATP concentration functions 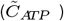, fitted signal functions (*χ*), and their product 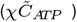. The glucose-restricted signal starts earlier and last longer than control, but has a lower peak amplitude. This likely occurred due to the release of additional reactive oxidative species during glucose-restriction. However, 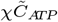 is diminished overall in glucose-restricted cells, signifying diminished contractility

As expected, glucose restriction caused the average ATP concentration to decrease, with glucose restricted cells starting at 46.5% of the control cell ATP content initially. Additionally, glucose restriction caused ATP to deplete instead of accumulate during contraction; glucose-restricted cell ATP decreased to 28% while control cell ATP increased to 116.5% at the end. Cells likely enrich themselves with ATP as a natural means of enhancing contractility (37). However, without sufficient glucose, this mechanism is impeded as anaerobic glycolysis fails to meet cellular demands, and ATP instead diminishes as contraction proceeds.

### Contraction Signal

Cell contraction is mediated by Rho GT-Pase through its downstream effector ROCK (Rho-associated-serine/threonine kinase), which phosphorylates myosin light chains and promotes the binding of myosin heads onto actin (38). The release of reactive oxidative species, such as that occurring under heightened UV exposure, is known to affect Rho signaling pathways and therefore tension generation in cells (39, 40). We assume this to be the dominant effect of UV-stimulation, and thus localize signaling (*χ*) entirely in cross-bridge kinetics (5).

The sigmoidal shape of our contraction signatures (Figure 2c) represents a gradual rise and subsequent decay in contractility, indicative of cooperative and competing effects in the dynamic process of cytoskeletal activation. To capture such effects in this context, we chose the standard auto-catalytic reaction rate function for our signal

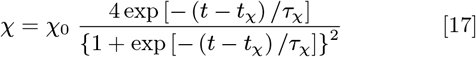

to model the UV-stimulated contraction signal, where *χ*_0_ is the signal amplitude, *t*_*χ*_ the transition time between rise and decay, and *τ*_*χ*_ the characteristic time controlling the kinetics of the process.

### Model Calibration

We use the three parameters of the signal function (*χ*_0_, *t*_*χ*_, *τ*_*χ*_) as calibration parameters for our model. Namely, we will look to fit our model against the normalized length contraction curves to derive the signal function for both the glucose-restricted and control cells. Ideally, we would fit individual length contraction curves with our model and derive signal functions for each cell. However, for simplicity, we fit the average length contraction curves (see colored *L/L*_0_ lines in Figure 2c), taking them as representative of the control and glucose-restricted cell population responses.

To perform the fits, we relate the lengthwise normalized contractions and our model variables

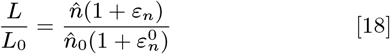

where 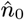 and 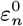 are the number of sarcomeric units in the SFs and their stretch at the start of contraction.

### Model Parameters

Next, we determine the values of the model parameters, compiling them in Table 1. From our experimental results (Figure 2), we determined that the control, glucose-restricted, and intermediate cells were initially equal in length on average (*L*_0_ ≈ 60 *µm*). We also set this length as our reference (*L*_0_ = *L*_*r*_). Furthermore, by assuming quiescent conditions prior to imaging, we set the SF strains to the steady-state value initially 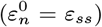. Thus, the initial number of sarcomeric units in all three cases must also be equal, which we set to be the reference number of sarcomeric units (*n*_0_ = *n*_*r*_). The stiffness of individual myosin necks was obtained from measurements on cross-bridges in muscles (*κ*_*m*_ = 1.75 *pN/nm*) (28, 41). To determine the characteristic neck stretch (*l*_*s*_ = 6.5 *nm*), we take the mechano-chemical efficiency of the cross-bridge cycle to equal the maximum measured sarcomere efficiency of 40% (42). Given ATP hydrolysis releases approximately 83 *zJ* of chemical energy (43, 44), the computed value of characteristic stretch imbues the myosin necks with a consistent amount of elastic energy in the attached state 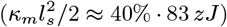. The value of the Cauchy parameter (*ε*_*s*_ = 0.225) was determined to qualitatively match the experimentally measured strain-dependence of sarcomere tension (31). We set the ATP parameters to match the averages reported in Figure 2d 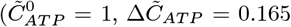 for control, and 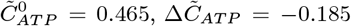 for glucose-restricted). Subsequently, we calibrate the myosin enthalpy (− (*µ*_*d*_ − *µ*_*a*_) = 0.362 *k*_*B*_*T*) to match the experimentally measured attached head fractions 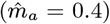 under regular isometric conditions 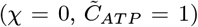 (41, 45, 46). For the coefficients of the SF configurational energy, and subsequently the steady-state stretch (*β* = 1.2, *p* = 2, *ε*_*ss*_ = 0.354), we adopted values used in the original SF kinetic model (47). The SF remodelling rate (*r*_*n*_ = 1*mHz*) was set to match values used by comparable models of SF kinetics (47–49). Lastly, to determine the normalized FA viscosity (SI: FA Kinetics)

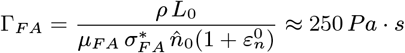

we take the linear coefficient for the tension-dependence of FA area 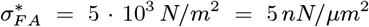, Eq.2) from measurements on 3T3 fibroblasts and MCF-7 (breast cancer) cells (20). The density of surface receptors in FAs (*ρ* = 270 10^12^ *m*^−2^ = 270 *µm*^−2^, *i*.*e*., spaced by roughly 60 *nm*) was estimated by averaging the values used by (50, 51) in their FA kinetic models, from which we also extracted the FA mobility (*µ*_*FA*_ = 5 *·* 10^4^ *m/Ns* = 50 *µm/nNs*) (50).

**Table 1.** Model parameter values.

| Symbol(s) | Value | Source |
| --- | --- | --- |
| $L_0$ | $60 \mu m$ | Figure 2b |
| $n_0, \varepsilon_n^0$ | $n_r, \varepsilon_{ss}$ | - |
| $\kappa_m$ | $1.75 pN/nm$ | (28, 41) |
| $l_s$ | $6.5 nm$ | (42–44) |
| $\varepsilon_s$ | $0.225$ | (31) |
| $-(\mu_d - \mu_a)$ | $0.362 k_B T$ | (41, 45, 46) |
| $\beta, p, \varepsilon_{ss}$ | $1.2, 2, 0.354$ | (29) |
| $r_n$ | $1 mHz$ | (48, 49) |
| $\mu_{FA}$ | $5 \cdot 10^4 m/Ns$ | (20, 50, 51) |
| $\tilde{C}_{ATP}^0, \Delta \tilde{C}_{ATP}$ | (ctrl.) $1, +0.165$ | Figure 3a |
| | (gl. res.) $0.465, -0.185$ | (averages) |

### Initial conditions

We assume that quiescent conditions prevail prior to imaging (at *t*_0_): without signaling (*χ* = 0), with the SF strain set to the steady-state value (*ε*_*n*_ = *ε*_*ss*_),and with the ATP concentration set to the initial value 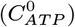.

Subsequently, we obtain the following expression for the initial SF tension

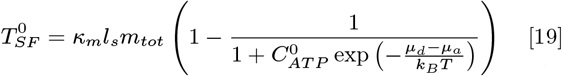

The fits were performed in MATLAB (R2021b) using their native least squares fitting algorithm (’lsqcurvefit’) with default fitting parameters. The same initial guesses for the signal parameters were used for the control and glucose-restricted fits, with results compiled in Table 2. In both cases, the fits successfully converged, yielding very good agreement with the experimental data (*R*^2^ = 0.998, 0.997), and thus, demonstrating that our model captured the contraction signatures well (Figure 3b).

**Table 2.** Fitted and initialized signal parameters.

| Case | $\chi_0$ | $t_\chi$ (s) | $\tau_\chi$ (s) | $R^2$ |
| --- | --- | --- | --- | --- |
| Control | 0.94 | 1970 | 222 | 0.998 |
| Glucose-restricted | 0.77 | 1800 | 408 | 0.997 |
| Initialization | 0.68 | 1997 | 392 | - |

## Model Results

### Fitted Results

Figure 3c plots the evolution of the fitted signal functions (1 + *χ*), the ATP concentrations 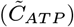, and their product 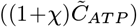. The control signal has a slightly higher peak amplitude (Δ*χ* = 0.94) compared to glucose-restricted (Δ*χ* = 0.77), with the control signal peaking slightly after the glucose restricted signal: *t*_*χ*_ ≈ 32 *min* for control, *t*_*χ*_ ≈ 30 *min* for glucose restricted. On the other hand, the characteristic time of the control signal (*τ*_*χ*_ = 222 *s*) is significantly smaller than glucose-restricted (*τ*_*χ*_ = 408 *s*), and consequently, the glucose-restricted signal develops earlier and decays slower than control. Glucose-restriction is also known to release reactive oxidative species in cells (52). This likely contributed to the oxidative species released through UV-stimulation, in turn, accelerating and prolonging the contractile signal. However, the product of the signal and ATP concentration 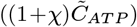 - which represents the effective factor that drives contractility - starts to develop at similar times for both cell populations (*t* ≈ 10 *min*), with the glucose-restricted value peaking slightly earlier, with significantly lower amplitude (both in absolute terms and relative to their starting values), and diminishing much sooner than control. Overall, this signifies that factors driving contractility are diminished as a result of glucose-restriction.

### Intermediate Glucose Restriction

We assume that the response to an intermediate level of glucose restriction lies between that exhibited by the control and glucose restricted cells. We assume that the intermediate response can be estimated by linearly interpolating the signal (*χ*) and ATP 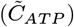 functions between the control and glucose-restricted values, based on the variable *ξ*, indicating the extent of glucose starvation (Figure 4a). Thus, *ξ* = 0 gives the control response, while *ξ* = 1 gives the glucose-restricted one. Because the extent of glucose restriction is given by glucose concentration *C*_*g*_, we define

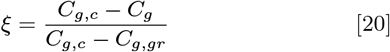

where *C*_*g,c*_ and *C*_*g,gr*_ are the glucose concentrations at control and (full) glucose restriction, respectively. Given this, in Figure 4b we compare the predicted and measured total contracted length (Δ*L/L*_0_) as a function of *ξ*. We see that the model prediction agrees well with the intermediate cell response (green datapoint, *ξ* = 0.76).

**Fig. 4.**
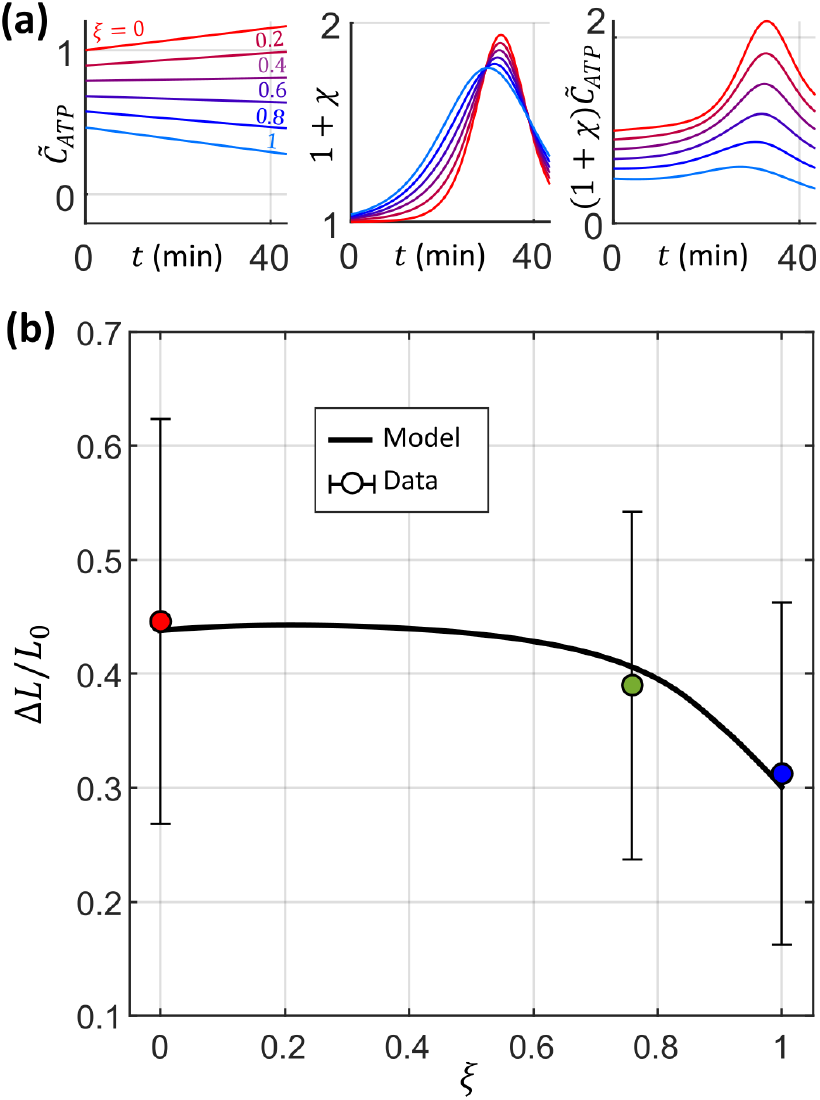
Results of interpolating between the response of the control and glucose-restricted cells. (a) The ATP concentration 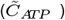, signal function (*χ*), and their product 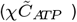 as a function of the interpolation parameter (*ξ*). (b) The total contracted length normalized by the initial cell length (Δ*L/L*_0_). Our model predicts a critical level of glucose restriction before contractility starts to significantly decrease relative to control (*ξ >* 0.45).

Our model predicts a small increase in total contracted length initially, reaching a maximum of 0.442 at *ξ* = 0.21. Then, the total contracted length decreases gradually with increasing *ξ*, only dropping below the control value (Δ*L/L*_0_ *<* 0.438) when *ξ >* 0.45. Further increasing *ξ* beyond this point is required before the total length contraction starts to diminish more significantly.

These results suggest that, subject to low levels of glucose-restriction (*ξ <* 0.21), cells adapt their metabolism to more efficiently use ATP and achieve slightly heightened levels of contraction. This parallels observations made on the metabolic adaptation of the human musculoskeletal system (53, 54). More importantly, however, it suggests that a critical level of glucose restriction must be exceeded before contractility becomes significantly affected. This may explain the inconsistencies in the response of glucose-restricted cells (13) in the literature. Below a critical severity - which likely depends on cancer phenotype (55) - glucose restriction is at best ineffective, and at worst detrimental, instead enhancing cell contractility and possibly advancing the rate of proliferation.

## Improvements to the Present Analysis

To further validate our analysis, additional experiments need to be conducted. Namely, more data can be collected at intermediate glucose restriction severities (Figure 4) to support our model prediction. Additionally, traction force microscopy and actin fluorescence data can be collected to validate the predicted SF tension 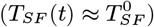 and SF shortening kinetics 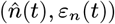. This will allow us to identify shortcomings in either the selected model parameters or the various simplifications in aspects such as signaling and FA kinetics.

## Discussion and Conclusions

We developed an experimental framework whereby glioma cancer cells undergo glucose restriction and are subsequently prompted to contract through UV stimulation. With a physics-based model, describing cytoskeletal contraction as a function of glucose availability, we predict changes in the contraction signature under various levels of glucose restriction. The model is based on an input signal for cytoskeletal activation, which we extrapolate from model calibration for (i) control and (ii) maximal glucose restriction, whereby we then estimate cell contraction at intermediate levels of glucose restriction and observe good experimental agreement. We observe that maximal glucose starvation effectively reduces cytoskeletal contraction, while intermediate levels of glucose starvation yield no significant reduction in contraction.

To better understand this outcome, we assumed that the response of cells subject to an intermediate level of glucose restriction may be interpolated from the control and glucose-restricted responses. Our model results suggest that cancer cells adapt their metabolism at low levels of glucose restriction, slightly improving their contractility overall. But the mechanisms for adaptation begin to fail above a critical level of glucose restriction, at which point contractility drastically diminishes. At sub-critical glucose restriction, our model predicts a slight increase in contraction. Our experiments, on the other hand, do not highlight an increased contractility but show negligible reduction. Our investigation, nevertheless, shows that low levels of glucose restriction are ineffective in reducing contractility, and therefore can be ineffective or possibly even detrimental in impeding cancer proliferation. This has widespread implications, given most glucose-restriction studies look at only a single severity. To better explore glucose restriction as a potential avenue for cancer therapeutics, it is imperative to conduct studies that sample various *levels* of glucose restriction to identify the critical conditions yielding reduced contractility in different cancer phenotypes. An important question to then ask is whether the critical glucose restriction level can be physiologically achieved in-vivo, and what are its general physiological implications.

## Materials and Methods

### Cell Culture

U-87 MG cells, derived from human glioblastoma-astrocytoma, were originally purchased from American Type Culture Collection (ATCC®HTB-14 TM) and maintained in our laboratories following ATCC protocols. Cells were grown in full media composed of high glucose (4.5 g/L) DMEM (STEMCELL Technologies) basal media supplemented with 10% FBS (Gibco, cat 12483020) and 1x GlutaMax (Thermo). Cells were subcultured at a 1:7 or 1:10 split ratio by passaging when 70-80% confluent, using 2.5 Trypsin-0.53 mM EDTA (1mL).

### Cell Preparation

Three 96 well-plates (Greiner *µ*clear, one with black walls for imaging and two with white walls for ATP measurements) were coated with Type I collagen (PureCol(r), Advanced Biomatrix) according to manufacturer’s protocol. Cells were seeded at a density of 3000 cells/well in full media and allowed to attach and grow over 72 hours in an incubator (37C, 5% CO2).

### Cytoplasmic Staining

Twenty-four hours prior to imaging, the cells’ cytoplasm was stained with Cell Tracker Green (Thermo), according to manufacturer’s protocol. A 10 *µ*M staining solution was prepared in FBS-free DMEM and added to cells after removing the full media from the wells. Cells were placed back in the incubator (37C, 5% CO2) for one hour, then the staining solution was washed and replaced again with full media. Cells were placed back in the incubator until the next day.

### Cell Conditioning (Glucose-restriction)

To condition the cells for imaging and ATP measurement, the media in each well was removed and replaced with conditioning media for two hours. glucose free DMEM (Thermo) without FBS and GlutaMAX was used for full glucose-restriction, glucose free DMEM supplemented with 10% FBS and 1x GlutaMAX was used for the intermediate glucose level. Full media was used as a control.

### Nuclear Staining

After conditioning, the media was replaced with Live Cell Imaging Solution (LCIS, Thermo) supplemented with FBS (2%) to which 8 µM of Hoechst 33342 dye (Thermo) was added to stain cell nuclei. Cells were incubated in LCIS for 5 minutes.

### Cell Imaging

Cells were imaged on a Cytiva GE IN Cell Analyzer 2200 with a 40x objective (N.A. 0.60, ELWD, collar correction, Nikon) under ambient conditions, using a solid-state illumination source and a cooled sCMOS camera. Images from two fluorescence channels (DAPI, FITC) and brightfield were collected in a time series at an interval of 45 s for a total of 60 time points. Exposure times for each channel were set at 500 ms for DAPI (Ex/Em 390/435 nm), 100 ms for FITC (Ex/Em 475/511 nm), and 100 ms for brightfield. Each resulting image was 2048 × 2048 pixels in size with a spatial resolution of 6.15 pixels/µm yielding an area of 333 × 333 µm. Two sets of images each for control and glucose-restricted cells were collected, totaling 6 sets. To avoid drifting in the focal plane, at each time point the image was automatically re-focused based on the first channel to be imaged (DAPI).

### ATP Measurements

Identically prepared plates of cells as described above were used to evaluate the total ATP associated with each cell-well population at the beginning and end of imaging. A spectrophotometric assay, CellTiter-Glo 2.0 (Promega) was used. The assay reagent consists of a lysis buffer and thermostable luciferase that generates a luminescent signal proportional to the amount of ATP present in the cell population. The plates are equilibrated to room temperature (30 minutes), and the Titer-Glo one-step reagent added to the wells in a 1:1 ratio with the culture media. The plates are then placed on a shaker for 2 minutes to lyse the cells. The lysed cells are left for 10 minutes for the luminescent signal to stabilize before luminescent readings are measured (Fluostar, BMG Labtech). ATP concentrations in wells were determined using a calibration curve (SI: Figure S1) produced using the same reagents and ATP standards (Promega). Three measurements each for control and glucose-restricted cells at the start and end of imaging were obtained, totaling 18 measurements.

### FBS Glucose Measurements

To estimate the typical glucose content in FBS, we assayed 3 different FBS batches using the Glucose Assay Kit (Sigma Aldrich, GAGO20), one batch that was heat inactivated (STEMCELL Technologies lot 2030348) and kept at 4oC for > 1 years; and two batches (STEMCELL Technologies lot 2472937RP and 2095193) that were stored frozen since aliquoted. The FBS was diluted 20x and passed through a 5kDa MWCO spin filter (Agilent 5185-5991) after the filter received a pre-wash with PBS to prime the polyethersulfone membrane material (to avoid potential losses). For each measurement, 50 µl of the filtered FBS was incubated with xxx µl of assay reagent. The reaction was then stopped with 100 µl of sulfuric acid, before proceeding with the assay. 6 measurements were made for each FBS batch, yielding average glucose content of 1.32, 1.06, and 0.89 g/L for the three batches respectively. The standard deviations are 0.06, 0.03, and 0.01 g/L respectively (Data provided in SI). For our analysis we elected to use the values from the second batch: 1.06 ± 0.03 g/L 1 g/L.

### Cell Segmentation

The cell images were analyzed with a custom process involving ImageJ (56) and MATLAB (2021b). Briefly, the images were stabilized (57) and cell-substrate boundaries were identified from cytoplasmic (FITC) images using 0.65 of the binary threshold value determined by Otsu’s method (58). Cell-cell boundaries were then identified with MATLAB’s watershed algorithm with the nuclear (DAPI) images acting as cell identifiers (SI: Figure S2). Cells that intersect the image boundaries were omitted, yielding 21 control and 15 glucose-restricted cells remaining for analysis.

### Cell Geometry During Contraction

With the segmented cell boundaries, we extracted the cells’ area *A*, length *L*, and width *w* at each time point *t*, with values at the start of imaging (*t* = 0) given by *A*_0_, *L*_0_ and *w*_0_ respectively. Let the column vectors ***x***(*t*), ***y***(*t*) contain paired *x* and *y* coordinates of all *N* (*t*) pixels within a cell’s boundary, relative to their instantaneous centroid. *A*(*t*) is converted directly from *N* (*t*) according to 6.15^2^ pixels/*µm*^2^ as per image specifications. Then, to determine *L* and *w*, we first compute the covariance matrix of each cell’s initial geometry

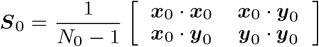

where ‘*·* ‘ is a dot-product operator and ***x***_0_ = ***x***(0), ***y***_0_ = ***y***(0). We define the first (***v***_*L*_) and second (***v***_*w*_) eigenvectors of ***S***_0_, as the respective directions of *L* and *w* for a given cell. Thus, we obtain *L*(*t*) and *w*(*t*) by computing the average absolute projections of ***x***(*t*), ***y***(*t*) onto these directions.

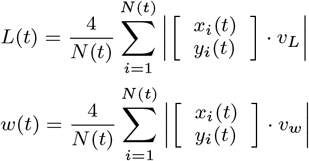

with *x*_*i*_(*t*),*y*_*i*_(*t*) the *i*^*th*^ entry in ***x***(*t*), ***y***(*t*), and the factor of 4 appropriately scales the computed averages to *L* and *w* (SI: Figure S3)

## ACKNOWLEDGMENTS

This work was supported by the New Frontiers in Research Funds – Exploration (NFRFE-2018-00730) and by the Natural Sciences and Engineering Research Council of Canada (NSERC) (RGPIN-2017-04464). We also thank Leonard Foster, Kyung-Mee Moon, and Vikram Deshpande for their help and consultation.

## Supporting Information for

### Supporting Information Text

#### Cell Contraction Energy Estimates

Table S1 reports first order estimates of the various energy dissipation terms associated with cell contraction. We find that dissipation associated with focal adhesions are greater than other terms by at least three orders of magnitude.

In these calculations we use length and velocity values that are representative of the observed cell contractions (see Figure 1b in the main text). Our cells are approximately 2000 *µm*^2^ in area (*A*), which equates to a circle of 50 *µm* diameter (*D*). Over a period of *t*_*im*_ = *t*_*f*_ − *t*_0_ = 45 *min* = 2700 *s*, they contract in their length by a characteristic strain of *ε* = 0.5, yielding a characteristic velocity of *v* = 10 *nm/s* and a characteristic strain rate of 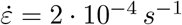. Idealizing the cell as a hemisphere, their characteristic volume computes to *V* 30 000 *µm*^3^.

Below, we describe in order, the equations used to estimate all the dissipation terms:

- **External hydraulic drag:** The drag force *F*_*d*_ on a sphere of radius *R* = *D/*2 = 25*·*10^−6^*m*, moving at a velocity *v* = 10^−8^*m/s* through a fluid of dynamic viscosity of *µ* = 1*·*10^−3^*Pa· · ·* (water) is given by Stokes’ law as *F*_*d*_ = 6*πµRv* = 4.71*·*10^−15^*N*. Half of this force (due to hemispherical shape) is multiplied by the sphere’s velocity to yield an estimate of the power done against external hydraulic drag, giving *p*_*hd*_ ∼ *F*_*d*_*v* = 4.71 *·* 10^−23^*W*.
- **Cytoplasmic viscosity:** A high estimate for cytoplasmic viscosity is *µ*_*cyt*_ = 1000 *Pa s* (1), which we multiply by the strain rate squared 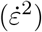 and integrate over the cell volume (*V*) to yield the rate of cytoplasmic viscous dissipation, giving 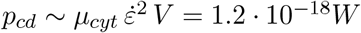.
- **Cytoskeletal permeability:** Flow of the cytosol through the permeable cytoskeleton dissipates energy. We implement the model described by (2). The force exchanged between the two is proportional to their relative velocity (*v*) and drag coefficient (*ξ*_*perm*_), which when multiplied again by the velocity yields a dissipative power density. This is (*φ*_*cs*_) and integrated over the cell’s volume (*V*) to yield the total dissipation due to cytoskeletal permeability.
- **Membrane transport:** There is dissipation associated with the transport of fluid through the cell membrane. The velocity of fluid *j*_*f*_ through the cell membrane is proportional to the membrane permeability *α*_*m*_ = 1 *·* 10^−4^*µm/P a s* (3) and the difference in chemical potential of fluid across the membrane Δ*ψ*_*f*_ ≈ 1000 *Pa*, obtained by assuming *j*_*f*_ = *v*. The dissipated power is given by integrating the flux over the total area *A* = 2*π R*^2^ = 3.93 *·* 10^−9^*m*^2^ and multiplying by the difference in chemical potential Δ*ψ*_*f*_, giving *p*_*md*_ ∼ Δ*ψ*_*f*_ *A j*_*f*_ = 3.93 *·* 10^−14^*W*.
- **Focal adhesion:** If we coarse grain focal adhesions as a distributed adhesion between the cell and the substrate, there is a dissipated energy proportional to the surface energy *θ*_*F A*_ and the newly created surface area ≈*A*_0_ = *πR*^2^ = 1.96 *·* 10^−9^*m*^2^. The dissipative power is obtained by dividing the energy *θ*_*F A*_*A*_0_ by the contraction time *t*_*im*_. We obtain *θ*_*F A*_ = 30 *·* 10^3^*J/m*^2^ (4) from measurements of mechanical work needed to detach single murine fibroblast L929 adhered to collagen-coated polystyrene. The dissipated power is then *p*_*F Ad*_ ∼ *θ*_*F A*_*A*_0_*/t*_*im*_ = 2.18 *·* 10^−8^*W*.

As demonstrated above, the energetic dissipation generated by FA remodeling is larger than all other dissipation by at least six orders of magnitude. This justifies our hypothesis of neglecting all dissipation other than FA remodeling. In the next section we elaborate the FA remodeling model we propose in this study.

#### FA Kinetics

We assume that during cell contraction the FA area is constant 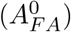 and set by the SF tension immediately prior to contraction 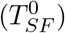

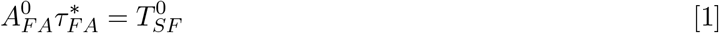

Thus, during sliding, the FA shear stress is related to the tension according to

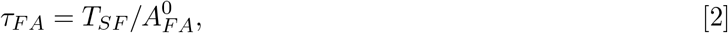

Above critical shear stress 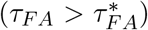, we assume that FAs slide at a rate inversely proportional to a viscosity *µ*_*F A*_ normalized by a characteristic length *l*_*s*_ and proportional to their stress (*τ*_*F A*_)

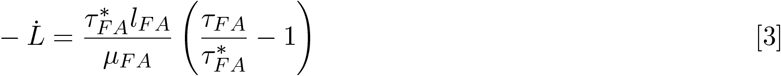

and, thanks to Eq. 1 and 2, Eq. 3 can be expressed in terms of the SF tension as

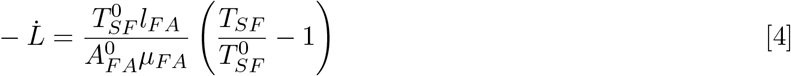

The rate of SF shortening 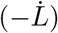 is related to the SF variables according to

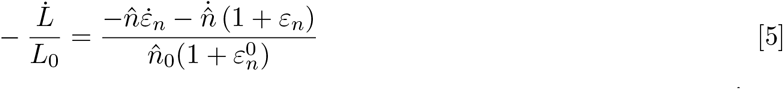

where *L*_0_ is the initial length of the SF, 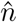, *ε*_*n*_ are their number of sarcomeric units and their stretch, 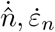 are their respective time-derivatives, and 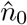, 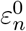 their respective initial values. Substituting Eq. 5, 2, and 1 nto Eq. 4 lets us derive the following governing equation for SF kinetics during FA sliding

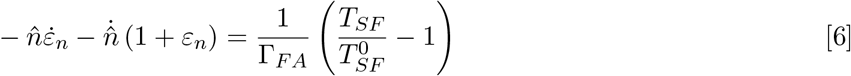

where Γ_*F A*_ is a new normalized viscosity

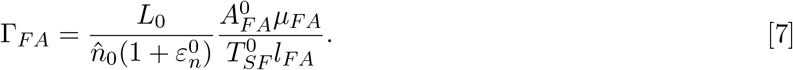

To determine an appropriate value for Γ_*F A*_, we express Eq.6 as

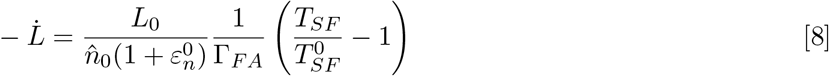

The FA kinetic model by (5) defines a characteristic mobility (*M*_*F A*_) to relate the sliding velocity of ndividual FA receptors - which we take to equal the velocity of the FA 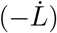, to the tension on individual eceptors (*T*_*r*_) - which we take relative to the receptor tension under quiescent conditions 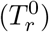

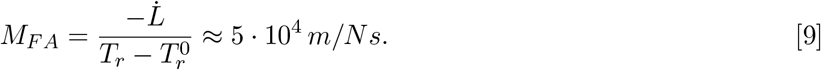

The tension in individual FA receptors is related to the total FA tension by

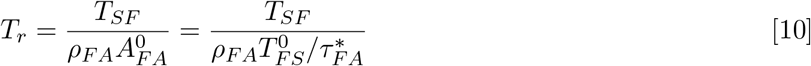

where *ρ*_*F A*_ (= 270 *·* 10^12^ *m*^−2^) is the density of receptors in the FA (5, 6), which we take to be a constant. The quiescent 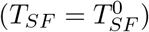 receptor tension is subsequently given by

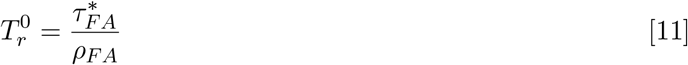

By solving for 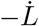 in Eq.9 and substituting Eq.10 and 11, we can compare the result with Eq.8, ultimately deriving the following relationship for the normalized FA viscosity

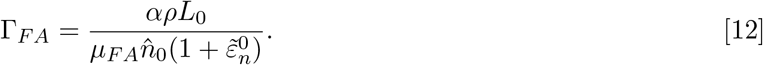

#### UV-stimulated Cell Contraction

In our study, cell contraction is triggered by exposure to ultraviolet (UV) and blue light. Prolonged and or intensified exposure to UV light induces irreversible changes in cells (phototoxicity), either directly by damaging DNA and by triggering apoptosis, or indirectly through the release of highly reactive oxidative species (7). UV exposure is typically minimized in experiments to better reproduce cell responses under natural conditions in-vivo. However, heightened UV exposure can also trigger cell contractility, namely, by altering contractile signaling pathways through the release of reactive oxidative species (8, 9). The triggered contractions typically complete within minutes to a few hours into UV exposure (10), and are additionally homogeneous and synchronous among cells - by virtue of the uniform illumination required for fluorescence imaging. By contrast, minimally motile cells need to be imaged for multiple hours and up to a day before significant motile response is observed, with the responses occurring without synchronicity and with substantial variability between cells (11). Triggering contraction by light exposure can therefore drastically shorten the duration of experiments. Furthermore, light-triggered contractility can be achieved very cheaply, requiring only the necessary equipment for fluorescent imaging. To capitalize on these benefits, we adopted UV and blue light exposure as a means of triggering controlled and coordinated contractions in our cells.

**Fig. S1.**
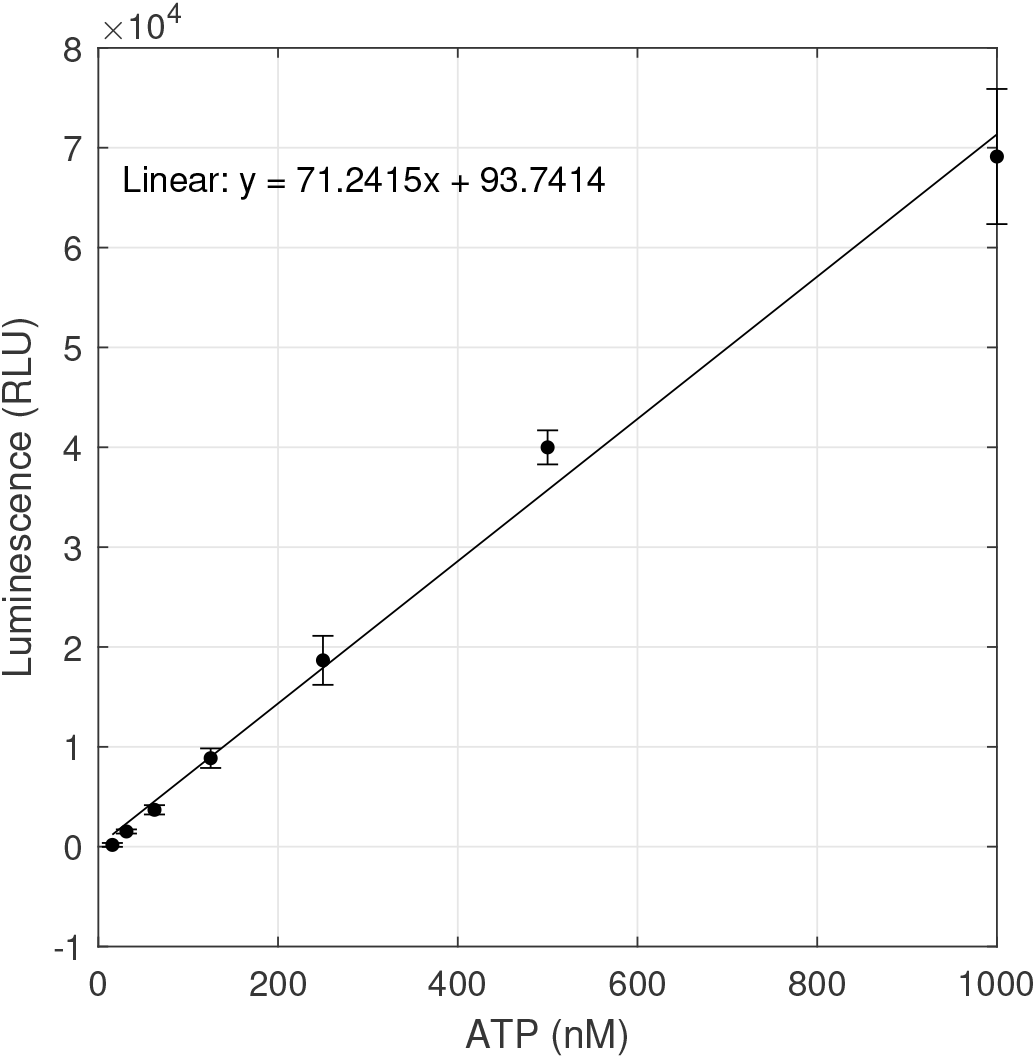
Calibration curve for ATP concentration data. Luminescence in relative light units (RLU) plotted against ATP concentration in nM. *R*^2^ = 0.993.

**Fig. S2.**
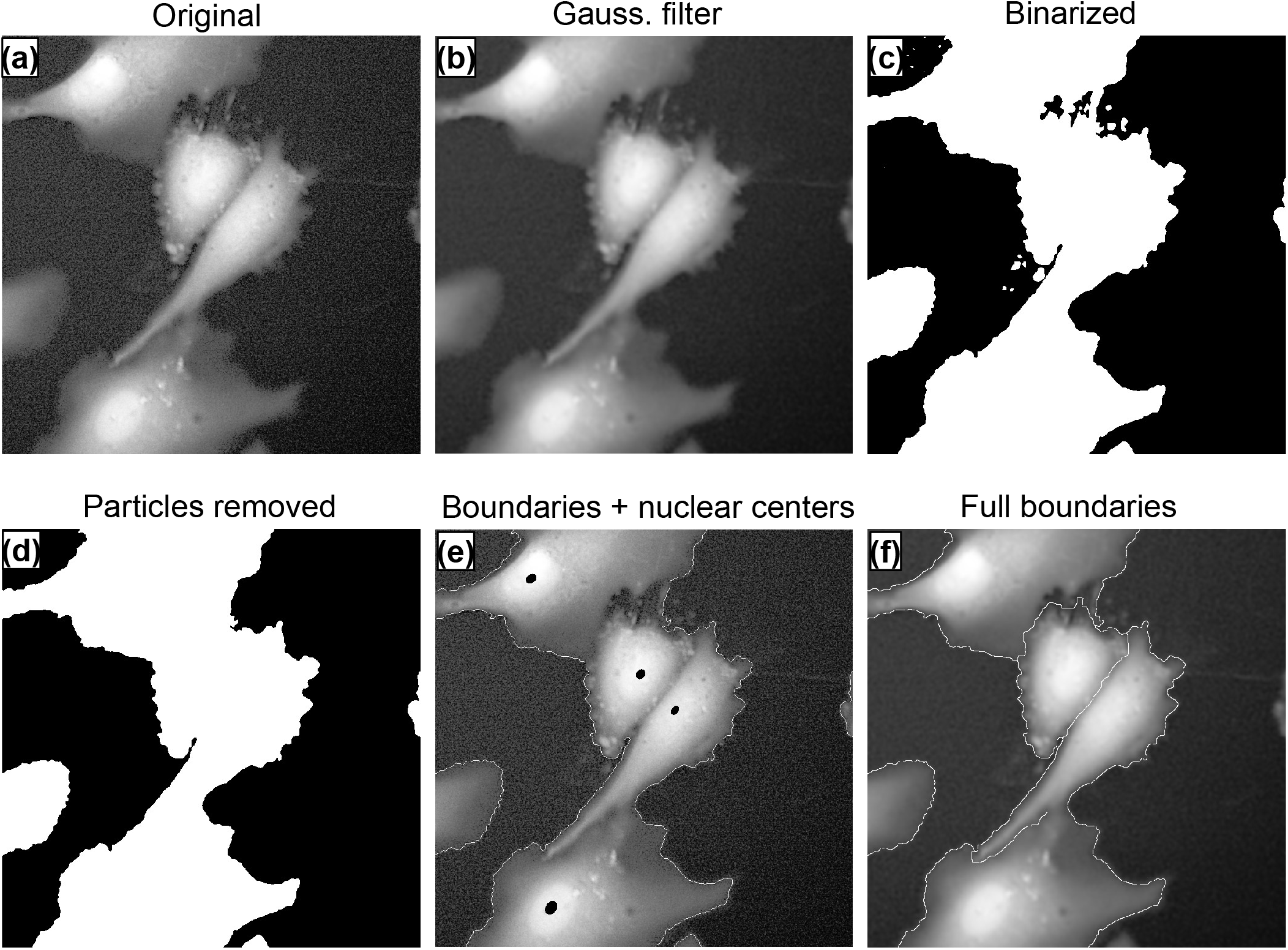
The image processing pipeline. (a) Histogram equalization (contrast adjustment) and stabilization in ImageJ. (b) Gaussian filtering in MATLAB with a 3 standard deviation smoothing kernel. (c) Binarization with 65% of the binary threshold value determined by Otsu’s method. (d) Holes and particles removed. (e) Identified cell-substrate boundaries and nuclear (DAPI) cell centers (black dot). (f) Full cell boundaries identified with MATLAB’s watershed algorithm.

**Fig. S3.**
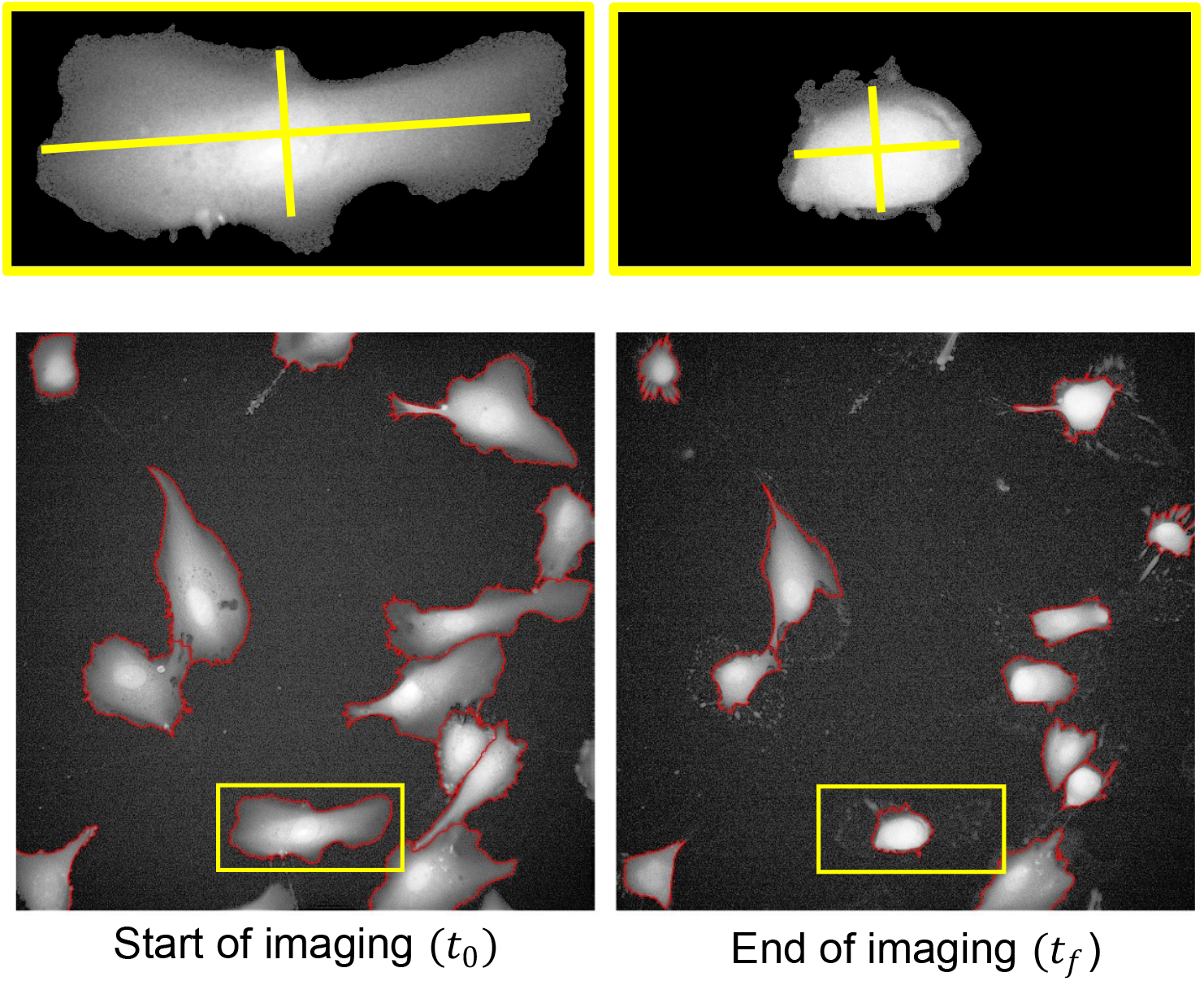
Sample computed cell length (longer yellow axis) and width (shorter yellow axis) for a control cell at the start and end of imaging. The orientation of the axes are maintained over time, but their centers are allowed to move, as described in the main text.

**Table S1.**
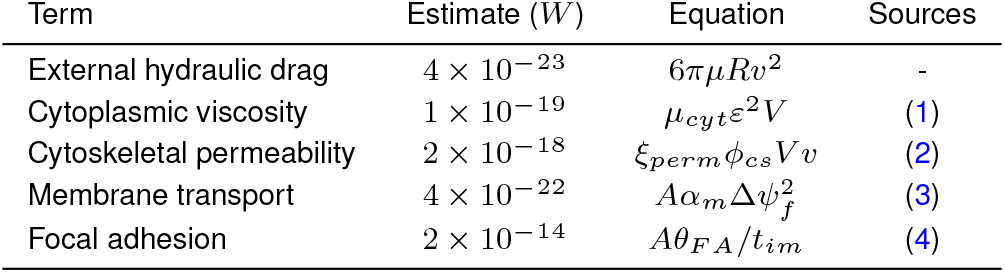
Estimates of dissipation terms associated with cell contraction.

**Movie S1. Video of the contracting cells with boundaries (red/blue lines). Top row = control, bottom row = glucose-restricted. Imaging time = 45 minutes**.

**Movie S2. Video of contracting cells that have been identified as suitable for data extraction (they do not intersect the video border and their boudaries were correctly identified). Separate colors were used to distinguish the cells. Top row = control, bottom row = glucose-restricted. Imaging time = 45 minutes**.

**SI Dataset S1 (GeomData.XLSX)**

Cell contraction geometry data (area/length/width for each cell considered in the analysis).

